# Rapid determination of the wide dynamic range of SARS-CoV-2 Spike T cell responses in whole blood of vaccinated and naturally infected

**DOI:** 10.1101/2021.06.29.450293

**Authors:** Anthony T Tan, Joey Ming Er Lim, Nina Le Bert, Kamini Kunasegaran, Adeline Chia, Martin Daniel Co Qui, Nicole Tan, Wan Ni Chia, Ruklanthi de Alwis, Ding Ying, Eng Eong Ooi, Lin-Fa Wang, Mark I-Cheng Chen, Barnaby Young, Li Yang Hsu, Jenny GH Low, David Chien Lye, Antonio Bertoletti

## Abstract

**Background:** Antibodies and T cells cooperate to control virus infections. The definition of the correlates of protection necessary to manage the COVID-19 pandemic, require both immune parameters but the complexity of traditional tests limits virus-specific T cell measurements.

**Methods:** We test the sensitivity and performance of a simple and rapid SARS-CoV-2 Spike-specific T cell test based on stimulation of whole blood with peptides covering the SARS-CoV-2 Spike protein followed by cytokine (IFN-γ, IL-2) measurement in different cohorts including BNT162b2 vaccinated (n=112; 201 samples), convalescent asymptomatic (n=62; 62 samples) and symptomatic (n=68; 115 samples) COVID-19 patients and SARS-CoV-1 convalescent individuals (n=12; 12 samples).

**Results:** The sensitivity of the rapid cytokine whole blood test equates traditional methods of T cell analysis (ELISPOT, Activation Induced Markers). Utilizing this test we observed that Spike-specific T cells in vaccinated preferentially target the S2 region of Spike and that their mean magnitude is similar between them and SARS-CoV-2 convalescents at 3 months after vaccine or virus priming respectively. However, a wide heterogeneity of Spike-specific T cell magnitude characterizes the individual responses irrespective of the time of analysis. No correlation between neutralizing antibody levels and Spike-specific T cell magnitude were found.

**Conclusions:** Rapid measurement of cytokine production in whole blood after peptide activation revealed a wide dynamic range of Spike-specific T cell response after vaccination that cannot be predicted from neutralizing antibody quantities. Both Spike-specific humoral and cellular immunity should be tested after vaccination to define the correlates of protection necessary to evaluate current vaccine strategies.

## Introduction

SARS coronavirus-2 (SARS-CoV-2), the etiological agent of coronavirus disease 2019 (COVID-19), has spread worldwide resulting in a global health and economic crisis that mass vaccinations are trying to resolve. The host’s capacity to be protected from viral infection or from the development of severe diseases requires a coordinated activation of different components of the immune system that ultimately lead to the production of neutralizing and antigen binding antibodies and antiviral T cells. Evidence that antibodies and T cells are required for protection have been generated in monkeys challenged with SARS-CoV-2 (1). Similarly, antibodies and T cells are present in the majority of SARS-CoV-2 infected individuals who control infection without severe symptoms (2–6) and a robust CD8 T cells response is associated with mild disease in oncological patients with humoral defects (7).

Recently developed SARS-CoV-2 vaccines that protect more than 90% of the vaccinated individuals from severe COVID-19 can induce Spike-specific antibodies and T cells (8, 9, 10). However, it is not fully clear which level of antibodies and/or T cells is necessary to exert such protection and whether differences do exist in antibody and T cell levels in vaccinated subjects. Efforts to define the protective threshold of antibodies through mathematical modelling (11) have shed some light on this issue but such work on T cell responses have so far been absent. While experimental data has shown that high levels of neutralizing antibodies can be sufficient to protect from experimental infection, lower levels require the presence of T cells (1). The level of neutralizing antibody titers are however extremely heterogeneous after natural infection(12) and while most of the new SARS-CoV-2 vaccines induce high neutralizing antibody levels (13), their persistence over time needs to be evaluated. Instead, virus-specific T cells appear to persist for a long time after viral clearance (i.e. 17 years after SARS-CoV-1 infection) and detection of SARS-CoV-2-specific T cells in COVID-19 patients with waning antibody titers have been reported by different groups (5, 4, 14, 15). Furthermore, the protective role of Spike-specific T cells in vaccinated individuals has also been highlighted by recent analysis of the early profile of Spike-specific immunity (16).

We think that the correlates of protection induced by vaccinations should therefore be derived from large prospective studies where both antibody and T cell levels are measured. However, while tests for antibodies are routinely performed, the technical complexity of SARS-CoV-2 T cell measurements have so far limited, with some exceptions (17), this analysis to few individuals characterized in few specialized laboratories. This is due to the fact that T cells specific for a defined pathogen are a minuscule fraction of the total T cells (often less than 1-3%) present in blood and can be distinguished mainly by complex functional assays that preserve the viability of the T cells during the assay. In addition, methods that are technically simple and do not require complex laboratory equipment, like ELISPOT, need to be performed in cells that have been purified from whole blood. This introduces into the assay the lengthy and technically demanding processes of peripheral blood mononuclear cell (PBMC) separation. Other assays that can directly measure the frequency and function of virus-specific T cells through expression of activation markers or cytokine production necessitate more complex equipment (i.e. Flow cytometer) and highly specialized personnel which might not be available in every routine diagnostic laboratory.

A possible rapid and simple alternative to these methods is the direct addition of stimulatory antigens or peptides to whole blood that results in the secretion of cytokines (usually IFN-γ) in plasma that is subsequently quantified. This assay is routinely applied for the diagnosis of active Tuberculosis (18) and it has also been shown to measure the presence of SARS-CoV-2-specific T cells in asymptomatic (5) and symptomatic SARS-CoV-2 infected patients (19, 20). However, to our knowledge, its accuracy and validation have not been properly analyzed over time in individuals who have been vaccinated against SARS-CoV-2, and only responses immediately after vaccination have been tested (16). Therefore, herein we utilized a range of cellular methods to measure SARS-CoV-2 T cell responses in individuals vaccinated with the prefusion stabilized, full-length SARS-CoV-2 S protein (BNT162b2) or naturally infected with SARS-CoV-2. We demonstrated that detection and relative quantification of Spike-specific T cells in vaccinated individuals can be easily and rapidly achieved through simple addition of Spike peptide pools to whole blood. Utilization of different peptide pools to stimulate whole blood provides the flexibility to derive rapid information about the kinetics and magnitude of Spike-specific T cell responses induced by vaccination and compare it with the one present in convalescent individuals.

## Results

### Rapid quantification of SARS-CoV-2 Spike-specific T cells by direct peptide stimulation of whole peripheral blood

We characterized the initial kinetics of Spike-specific T cells induced by two doses of mRNA vaccine (BNT162b2) over a 51 day period with different methods of antigen-specific T cell analysis in fresh blood and in cryopreserved PBMC. Whole blood of 6 healthy individuals was collected before (Day 0) and 7, 10 and 20 days after the prime and 7, 10, 20 and 30 days after the boost dose. Whole fresh blood (2-6 hours from collection) was either directly stimulated with peptides for a cytokine release assay (CRA) or processed to isolated PBMCs by Ficoll density gradient centrifugation (Figure 1A). PBMCs were either used fresh in an IFN-γ ELISPOT assay (Figure 1A) or cryopreserved for further analysis. Fresh blood and fresh PBMCs were stimulated with the SpG peptide pool containing 55 15-mer peptides (Supplementary Table 1) covering Spike-specific T cell epitopes that are immunogenic in 95% of SARS-CoV-2 infected individuals(5). Two negative controls consisting of the vehicle control with identical DMSO concentration present in the SpG peptide pool and a peptide pool covering SARS-CoV-2 nucleoprotein (NP; 41 peptides covering the C-terminal half of nucleoprotein; Supplementary Table 1) were used. Levels of IFN-γ and IL-2 were measured in the whole blood after 14-18 hours of incubation while the numbers of spots were also enumerated in the ELISPOT assay after overnight incubation.

**Figure 1.**
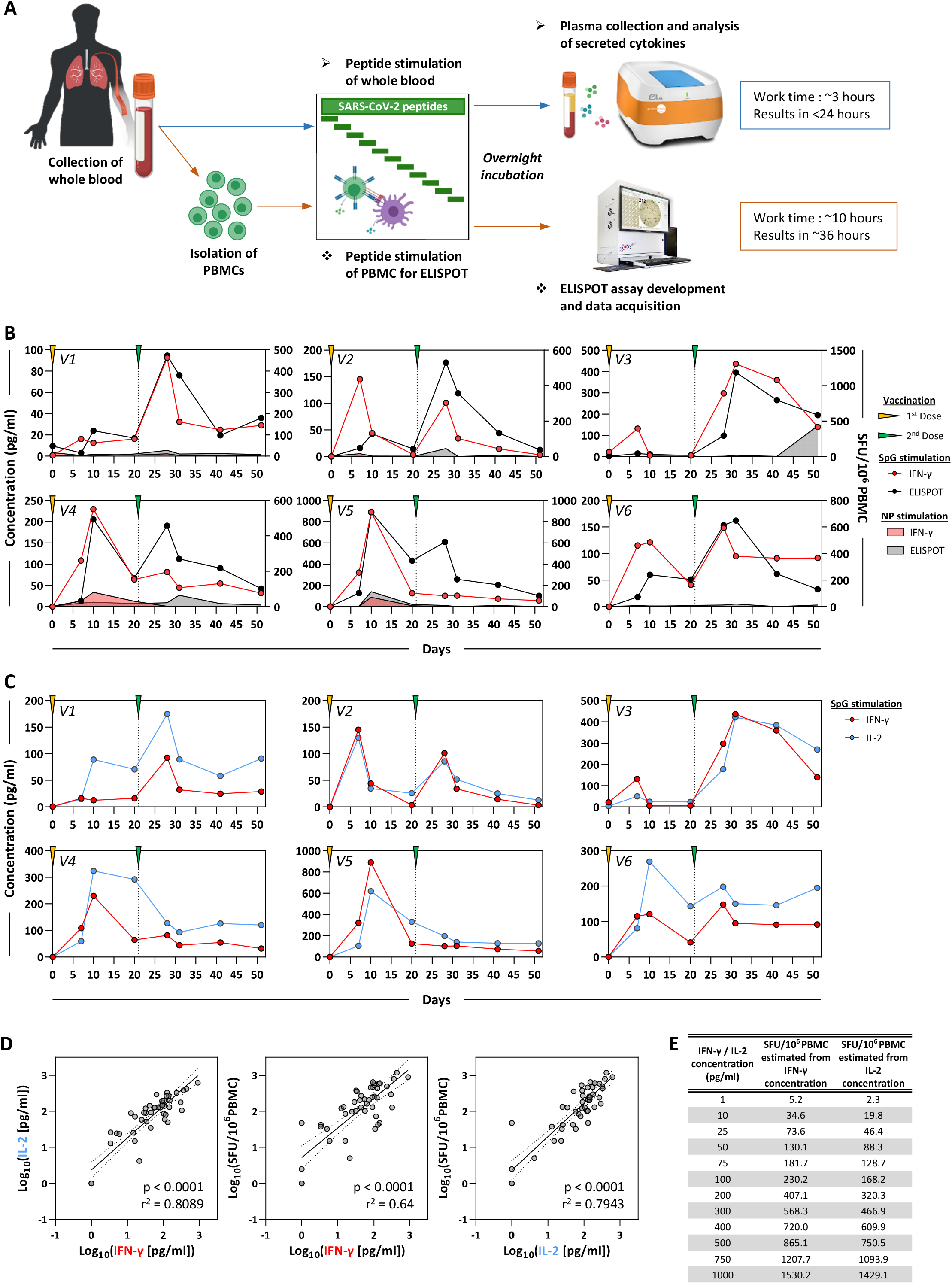
Detection of SARS-CoV-2 Spike-specific T cells by peptide stimulation of whole peripheral blood from vaccinated individuals. **A)** Comparative schematic representation of the workflow for the direct peptide stimulation of whole peripheral blood and the subsequent detection of cytokine secretion, and a standard IFN-γ ELISPOT assay. **B)** 6 healthy individuals were vaccinated with 2 doses of BNT162b2 according to the recommended schedule (21 days apart) and whole blood samples were longitudinally analyzed at 7, 10, 20 and 30 days after each dose. The collected whole blood was either directly stimulated for 16hrs with peptide pools specific for the Spike (line) or NP protein (shaded area), or immediately processed with Ficoll-density gradient centrifugation to isolate PBMCs. A standard IFN-γ ELISPOT assay using the SpG- or NP-specific peptide pools was then setup using the freshly isolated PBMCs. The quantity of secreted IFN-γ in stimulated whole blood (red line) was compared to the frequency of peptide reactive PBMCs quantified by IFN-γ ELISPOT (black line). **C)** The quantity of secreted IL-2 (blue line) in whole blood stimulated with SpG peptide pool are compared to the amount of IFN-γ detected. **D)** Linear regression analysis of the concentration of IFN-γ and IL-2 in SpG-specific peptide pool stimulated whole blood and the corresponding frequency of Spike-specific PBMCs (n=6; 48 samples). Dotted lines denote the 95% confidence interval. **E)** Table shows the estimated IFN-γ SFU/10^6^ PBMCs derived from the concentrations of IFN-γ and IL-2 in SpG peptide pool stimulated whole blood based on the linear regression analysis in D.

Figure 1B shows that the two different assays detected a predominant Spike-specific response in all the individuals and defined a matching profile of Spike-specific T cell response after the prime and boost vaccination doses. The number of IFN-γ spots detected after boost vaccination matches that observed in the phase 1/2 trial of individuals vaccinated with BNT162b2 (13) and with a similar preparation consisting of the trimerized secreted version of the Spike receptor-binding domain (BNT162b1) (10), including a trial conducted in Chinese individuals vaccinated with BNT162b1 (21). While stimulation with the NP-specific peptide pool remained largely negative, the IFN-γ quantities in blood and the number of IFN-γ spots showed identical peak responses that occurred 7-10 days after the first dose in individuals V4 and V5, and 7-10 days after the second dose in individuals V1, V3 and V6. There were however some minor discrepancies. CRA did not detect the boost of Spike-specific T cells induced by the second vaccine dose in subjects V4 and V5, perhaps in relation to the transient lymphopenia induced by mRNA vaccination (13). IL-2 cytokine measurement (Figure 1C) depicted an equivalent pattern of Spike-specific T cell response with that obtained through IFN-γ release. However, IL-2 levels exceed the level of IFN-γ in all the individuals 21 days after the first and second dose of vaccination. Overall a very strong correlation between IL-2 and IFN-γ secretion with the number of IFN-γ spots was evident (Figure 1D), which allowed a precise estimation of the quantity of IFN-γ producing cells related to the quantity of cytokines detected in whole blood (Figure 1E).

### Assessing the Spike-specific T cell response directly from fresh whole blood yields comparable results to classical T cell assays

Since T cell analysis is often performed in a single centralized laboratory using cryopreserved samples collected at different sites, we also analyzed the Spike-specific T cell response after vaccination using ELISPOT and activation induced cellular markers (AIM) assay performed using cryopreserved samples stimulated with SpG peptide pool and compared the results obtained with that from ELISPOT and CRA performed using the corresponding fresh whole blood. As already shown (22), the quantity of Spike-specific spots detected by ELISPOT in cryopreserved PBMCs was reduced in comparison to the ones detected in freshly isolated PBMCs (Supplementary Figure 1A), but the dynamics of the Spike-specific response remained consistent with fresh PBMCs (Supplementary Figure 1A) as also evidenced from the high correlation between the ELISPOT results from the differently processed samples (Figure 2A; Supplementary Figure 1B). The AIM assay in our hands was less precise in detecting the dynamic expansion and contraction pattern of the Spike-specific T cell response (Supplementary Figure 1C and D) likely due to the negative impact of cryopreserving PBMCs. Nevertheless, the AIM ability to differentiate between CD4 and CD8 T cell response represent an asset that should not be discounted.

We then assessed whether whole blood CRA results could reflect the results obtained using other assays (Figure 2A and B). We correlated the results obtained in all the different assays of Spike-specific T cells in cryopreserved and fresh PBMC samples with the results of the whole blood CRA. Cytokines in whole blood remained well correlated with ELISPOT assays that used cryopreserved or fresh PBMC samples, while its correlation with AIM assay results were generally weaker (Figure 2A and B). These results indicate that the CRA which utilizes freshly collected whole blood is a robust method that can reliably quantify Spike-specific T cell responses with results that are comparable to well-established assays used to analyze T cell responses.

**Figure 2.**
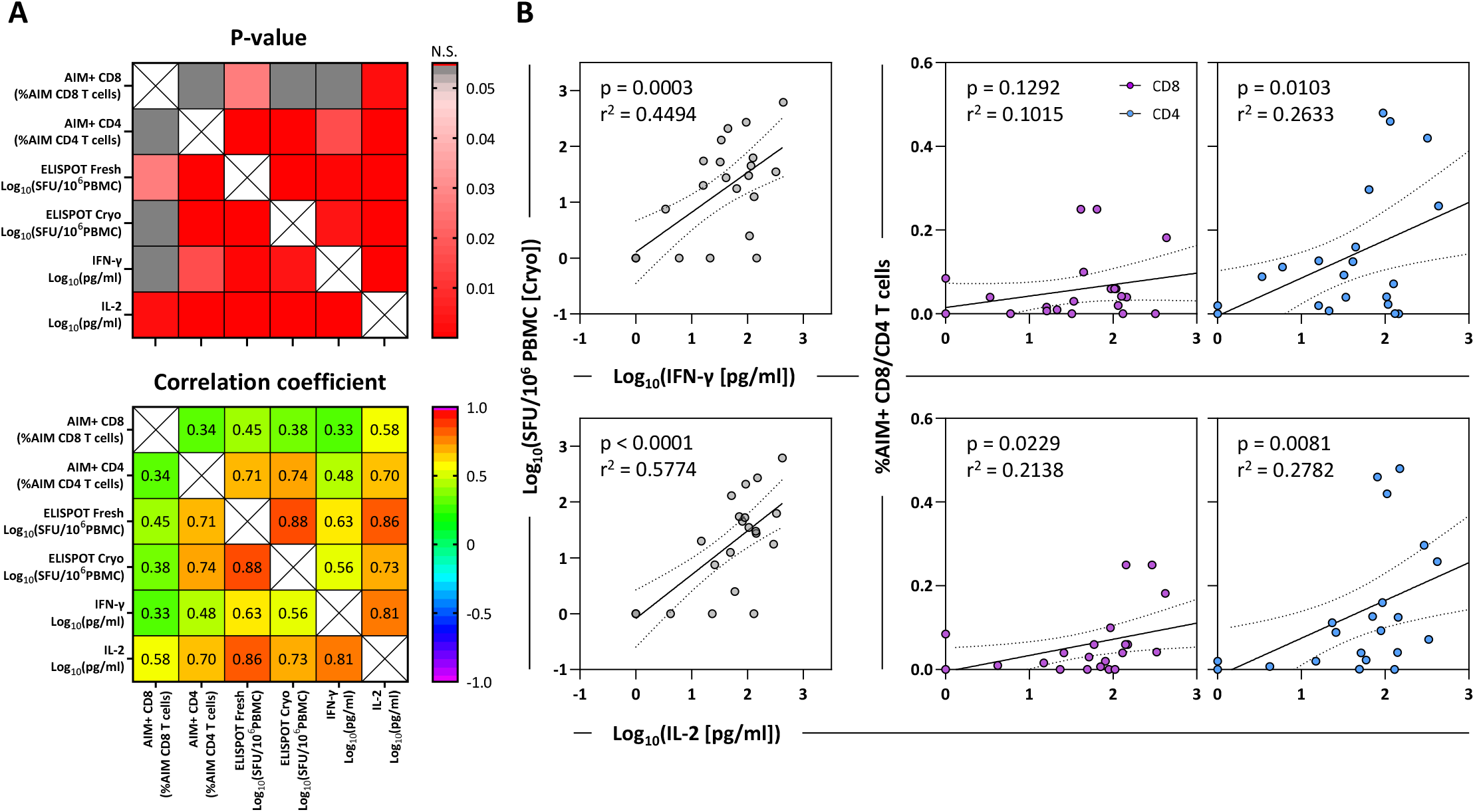
Correlation matrix of different assays used to quantify Spike-specific T cells. **A)** The top matrix denotes the significance of the correlation while the Spearman correlation coefficient is shown in the matrix below (n=6; 24 samples). **B)** Linear regression analysis of the concentration of IFN-γ and IL-2 in SpG peptide pool stimulated whole blood and the corresponding frequency of SpG reactive T cells in cryopreserved PBMCs quantified by either IFN-γ ELISPOT or AIM assay (n=6; 24 samples). Dotted lines denote the 95% confidence interval.

### Determining fine specificity of Spike-specific T cell responses through whole blood CRA

We next tested whether the whole blood CRA can be used to rapidly define T cell immunogenic regions of the whole Spike protein in vaccinated individuals. 15-mer peptides with an overlap of 10 AA covering the whole 1273 AA long Spike protein were organized in distinct pools of ~40 peptides that cover the following 7 Spike regions: Pool 1 (1-180), Pool 2 (171-345), Pool 3 (336-510), Pool 4 (501-705), Pool 5 (696-895), Pool 6 (886-1085), Pool 7 (1076-1273). A schematic representation of the localization of peptide pools 1 to 7 in relation to the S1 (N-terminal), RBD and S2 (C-terminal) regions of Spike is displayed in Figure 3A. These different Spike peptide pools were used in the whole blood CRA and ELISPOT with freshly isolated PBMCs. The results of these different assays performed at the indicated time points is first shown in two representative vaccine recipients (Figure 3B) while the results obtained in all the 6 different individuals are represented as a heat map displayed in Figure 3C. The three different measurements (IFN-γ and IL-2 CRA, and IFN-γ ELISPOT) provided very similar information in relation to the T cell response induced by BNT162b2 in healthy individuals. Even though some differences can be noted, like in individual V3, in whom the dominant IFN-γ response (CRA and ELISPOT) was induced by Pool 3 while Pool 7 induced the dominant IL-2 response, overall all the assays were largely equivalent. Consistent across the three different measurements, Spike-specific T cells preferentially targeted the S2 chain of Spike (covered by Pools 5, 6, and 7 spanning Spike 700-1273 AA) with responses in all the 6 individuals tested at different time points. Whole blood CRA and ELISPOT also showed that the region 501-705 AA contained in Pool 4 is the least immunogenic with only 1 out of the 6 vaccine recipients displaying a response at different time points (Figure 3B and C). Interestingly, analysis of individuals who recovered from SARS-CoV-2 infection (23) and in mRNA vaccine recipients (24) showed a similar reduced frequency of response in the Spike region 500-700 AA for CD4 T cells assayed through AIM detection. Taken together, these data show that direct analysis of cytokines secreted in whole blood pulsed with different peptides constitute a reliable method to gauge the presence and magnitude of functional T cells specific for epitopes covered by the utilized peptides.

**Figure 3.**
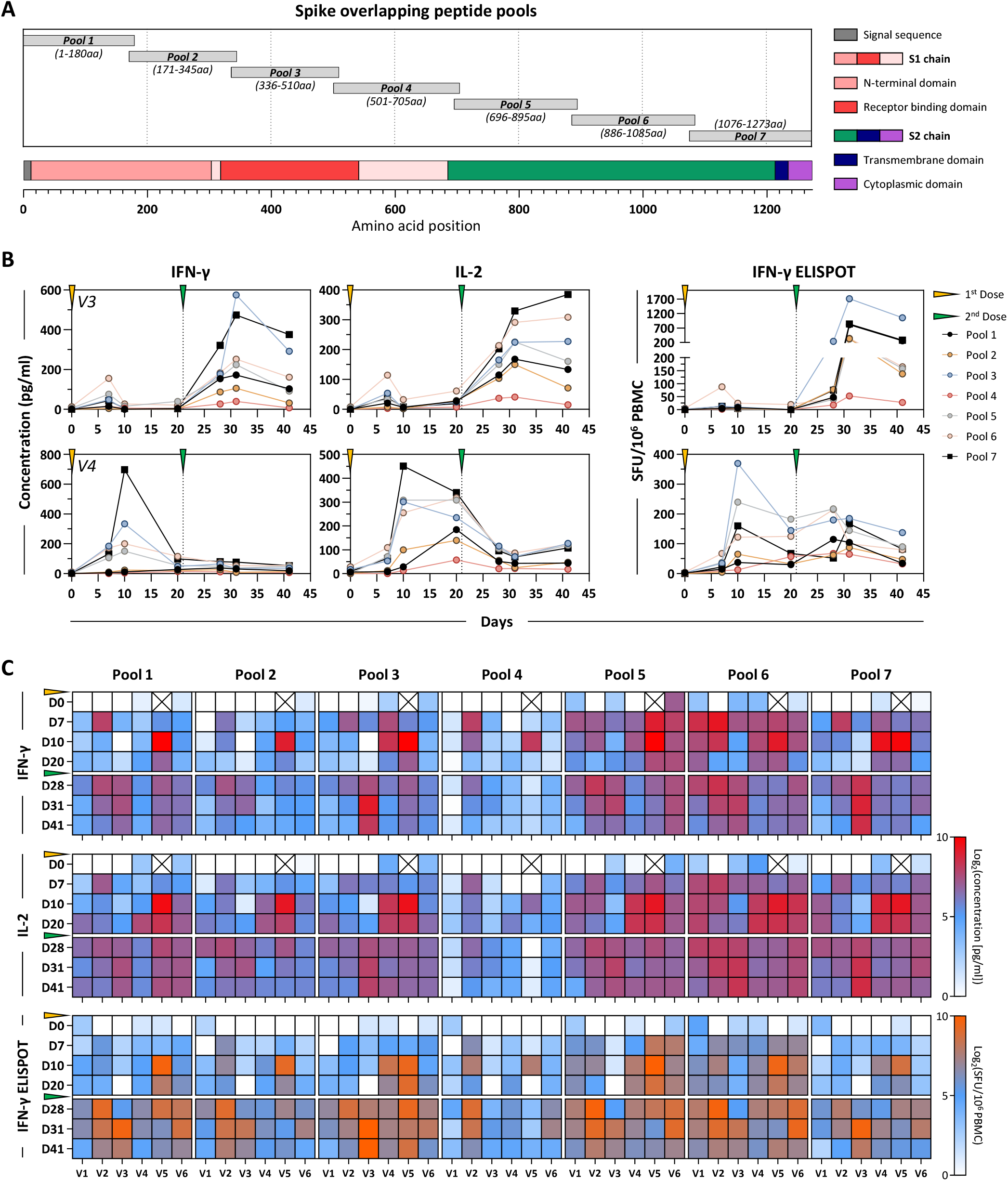
Immunodominance of Spike-specific T cells in vaccinated individuals. **A)** Schematic representation of the 7 Spike-specific peptide pools containing 15-mer overlapping peptides spanning the entire Spike protein. Pools 1-4 contain peptides from the signal peptide and the S1 chain while pools 5-6 encompass the S2 chain together with the transmembrane and cytoplasmic domains. **B)** Plots show the longitudinal evaluation of Spike-specific T cell responses (pools 1-7) through the quantification of IFN-γ (left) or IL-2 (middle) in peptide stimulated whole blood, or through IFN-γ ELISPOT (right) in 2 representative vaccinees. **C)** Heatmap shows the Spike-specific T cell responses quantified longitudinally in all vaccinees (n=6) using the 3 different assays described above. “X” denotes timepoints that were untested.

### Total Spike-specific T cell response is accurately represented by the T cells specific for the SpG peptide pool

While whole blood CRA using the 7 overlapping peptide pools of the Spike protein could give us information on the immunogenicity of the different regions of Spike and the total Spike-specific T cell response, the feasibility of assessing the response in larger numbers of individuals requires a more streamlined approach. As such, we analyzed the relation between the total Spike-specific T cell response and the response against our selected SpG peptide pool. A schematic representation of the localization of the peptides contained in the SpG peptide pool is displayed in Figure 4A. By correlating the results obtained from 3 different assays (IFN-γ and IL-2 CRA, and IFN-γ ELISPOT) where both the total Spike protein and SpG peptide pool-specific T cell response was determined in the same sample through stimulation with the corresponding peptide pools, we observed a strong positive linear relationship indicating that the T cell response against the SpG peptide pool is highly representative of the total Spike T cell response (Figure 4B). In fact, the SpG peptide pool-specific T cell response constitutes ~60-80% of the total T cell response against the entire Spike protein (Figure 4C). Hence, we proceeded to analyze the Spike-specific T cell response in a larger cohort of BNT162b2 vaccinated individuals and in individuals who have recovered from SARS-CoV-2 and SARS-CoV-1 infection using the whole blood CRA with SpG peptide pool as a stimulant.

**Figure 4.**
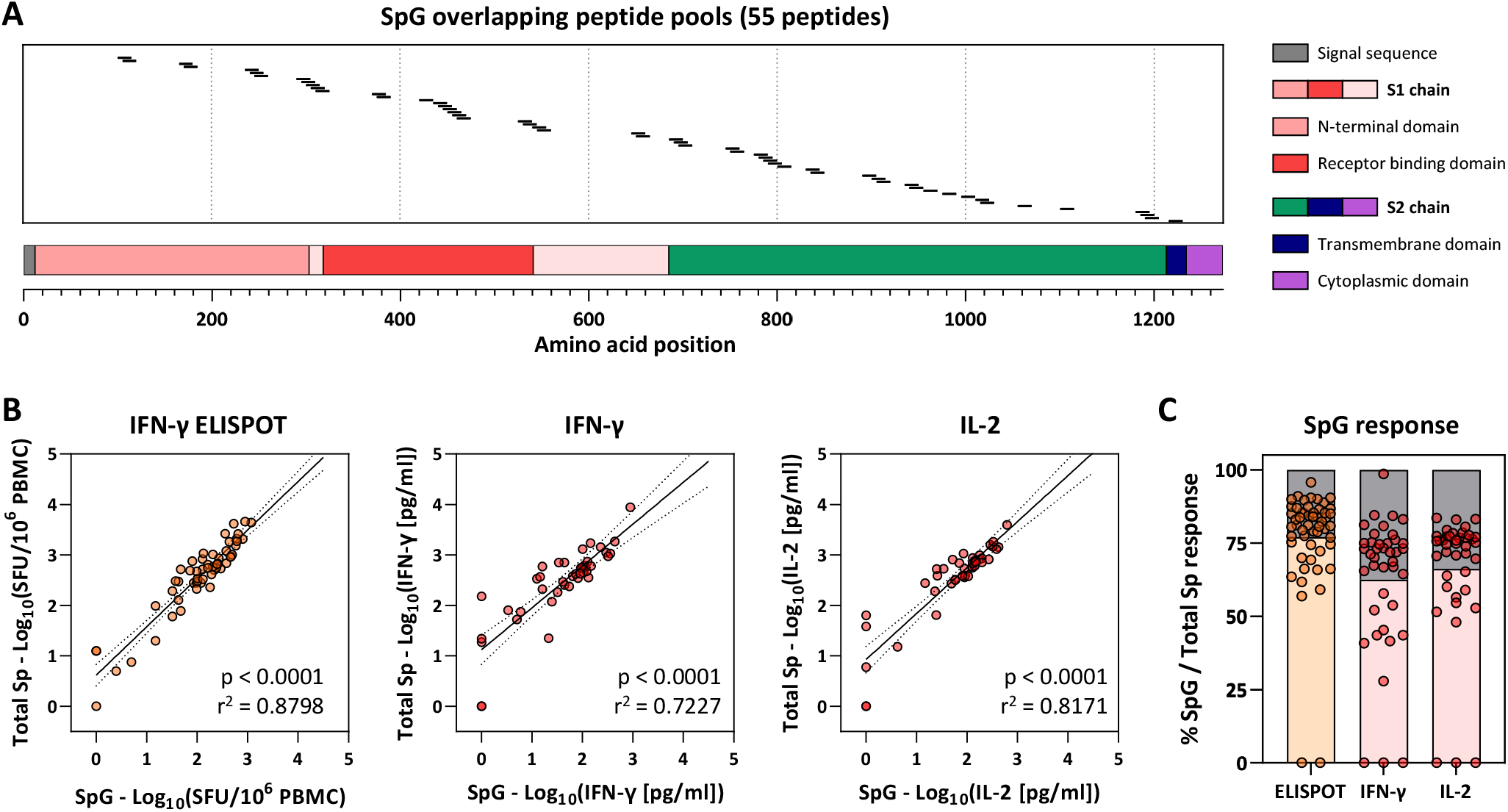
Frequency of SpG peptide pool and total spike protein-specific T cells. **A)** Schematic representation of the individual 15-mer overlapping peptides contained in the SpG peptide pool. **B)** Linear regression analysis of the T cell response against SpG peptide pool and the total spike protein (pool 1-7) as evaluated by ELISPOT (left) or by the quantification of IFN-γ (middle) or IL-2 (right) in peptide stimulated whole blood (n=6; 42 samples). **C)** The SpG peptide pool –specific T cell response quantified by each assay was expressed as a fraction of the total spike protein T cell response observed (n=6; 42 samples).

### T cell response to Spike after vaccination or after natural infection with SARS-CoV-2 or SARS-CoV-1

A total of 112 BNT162b2 vaccinated individuals (201 samples), 62 and 68 individuals who recovered from symptomatic (115 samples) and asymptomatic (62 samples) SARS-CoV-2 infection respectively, and 12 individuals who recovered from SARS-CoV-1 infection 18 years ago (12 samples) were studied longitudinally using the whole blood CRA with SpG peptide pool and measuring both IFN-γ and IL-2.

Firstly, we analyzed samples collected ≥3 months post boost vaccination (3 months) or SARS-CoV-2 infection clearance (3-12 months) to understand if the whole blood CRA remains reliable in quantifying the Spike-specific T cell responses at later time points beyond that tracked in Figure 1 (Supplementary Figure 3). Linear regression analysis of IFN-γ and IL-2 secretion from whole blood CRA and the corresponding frequency of IFN-γ spots in ELISPOT showed a good correlation across all subject groups at time points 3 months and beyond (Supplementary Figure 3). This indicates that whole blood CRA can quantify Spike-specific T cell responses accurately in vaccinated and infected individuals within 3 and 12 months after boost vaccination or infection resolution respectively.

Analyzing all the time points studied, we observed that vaccinated individuals exhibited a pronounced Spike-specific T cell response at the earliest analyzed timepoint (~14 days) after the boost vaccination dose which gradually declined and started to stabilize above the positivity threshold at 2-3 months (34/35 IFN-γ positive at 3 months), consistent with an initial T cell clonal expansion induced by vaccination and subsequent normalization (Figure 5A; Supplementary Figure 4). Similar kinetics were observed in natural SARS-CoV-2 infection where Spike-specific T cell responses were high around 1 month after infection clearance which gradually declined and stabilized above the positivity threshold (Asymp.: 23/27 positive; Symp.: 46/55 IFN-γ positive at 9-12 months) regardless of the symptomatic presentation during infection (Figure 5A; Supplementary Figure 4). Even in individuals who recovered from SARS-CoV-1 infection 17 years ago, T cells specific for SARS-CoV-2 Spike protein also remained detectable (8/12 IFN-γ positive) similar to the nucleoprotein-specific T cells described previously (25), despite the low AA conservation of the SpG peptides between the two viruses (Figure 5A). Clearly, whether T cells induced by vaccines will be maintained as the ones induced by natural infection beyond the 3 months period will have to be analyzed.

**Figure 5.**
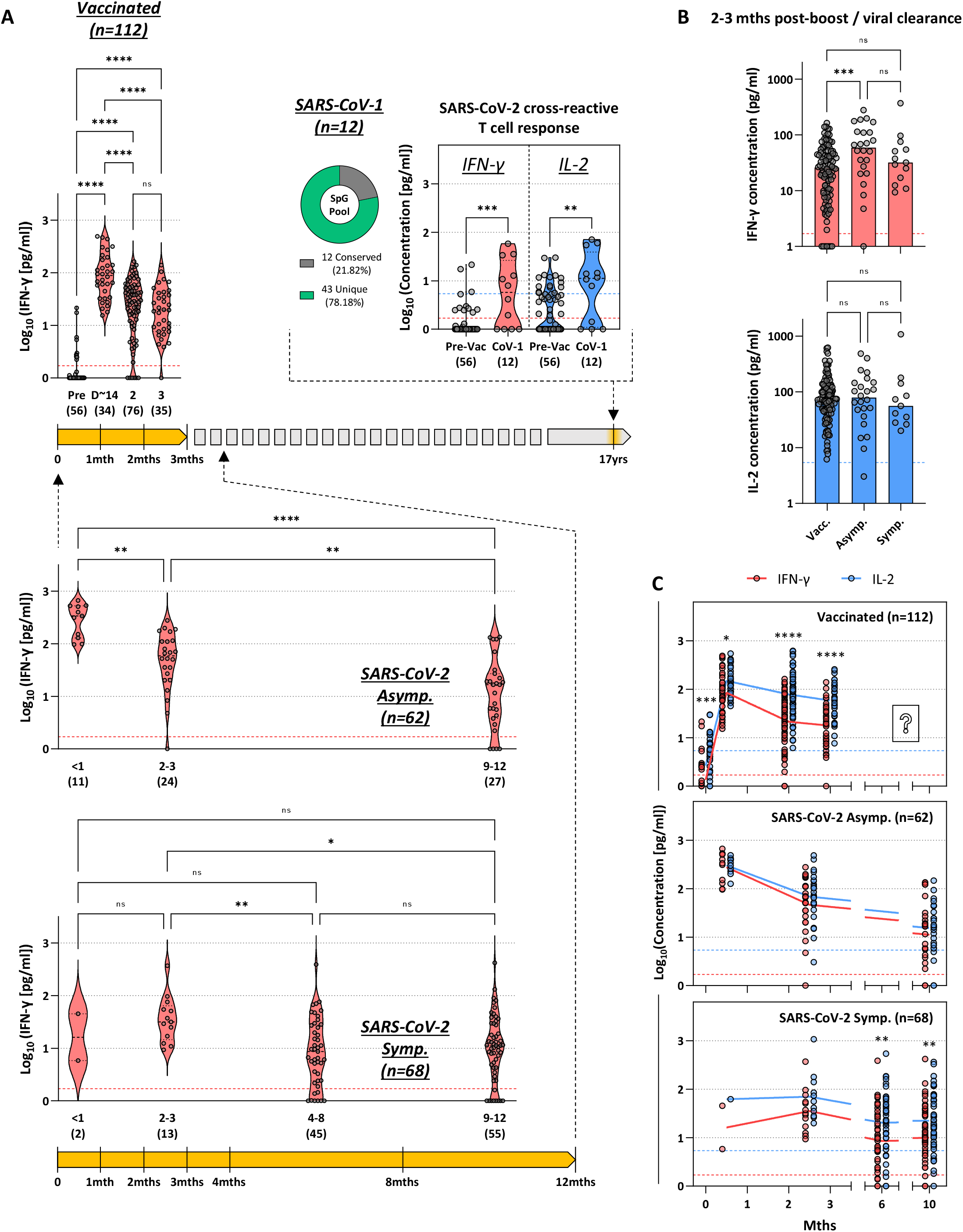
Longitudinal quantification of Spike-specific T cells from whole blood in vaccinated and infected individuals. **A)** SARS-CoV-2 Spike-specific T cell response in vaccinated individuals (n=112; 201 samples), convalescent asymptomatic (n=62; 62 samples) and symptomatic (n=68; 115 samples) COVID-19 patients were longitudinally quantified by measuring IFN-γ secretion in whole blood after SpG peptide pool stimulation. Cross-reactive SARS-CoV-2 Spike-specific T cells were also quantified in the whole blood of individuals who were infected with SARS-CoV-1 18 years ago (n=12; 12 samples). The response of individuals before receiving BNT162b2 vaccination are shown for reference. Pie chart denotes the number of peptides in the SpG peptide pool that are conserved or unique between SARS-CoV-2 and SARS-CoV-1. The sampling timespan (highlighted in yellow) is shown and the number of samples analysed at each time point were indicated in parentheses. Dashed lines denote the detection cut-off for the measured cytokines. Significant differences in each group were analysed by one-way ANOVA and the adjusted p-value (adjusted for multiple comparison) are shown. ns = not significant P>0.05; * = P≤0.05; ** = P≤0.01; *** = P≤0.001; **** = P≤0.0001. **B)** The quantities of secreted IFN-γ (red) and IL-2 (blue) in SpG peptide pool stimulated whole blood of vaccinees and COVID-19 patients sampled 2-3 months post-boost vaccination dose or viral clearance are shown. Bars denote the median value of each group and the dashed lines denote the detection cut-off for the measured cytokines. Significant differences were analysed and displayed as above. **C)** Longitudinal dynamics of secreted IFN-γ (red) and IL-2 (blue) in SpG peptide pool stimulated whole blood of vaccinees and COVID-19 patients. Significant differences were analysed and displayed as above. Dashed lines denote the detection cut-off for the measured cytokines.

We also assessed whether differences in the magnitude of the response were present between the different groups. We compared the Spike-specific T cell response detected at similar time points 2-3 months post vaccine boost or viral clearance. Unlike reports of higher quantities of neutralizing antibodies in vaccinees (13), individuals with symptomatic SARS-CoV-2 infection and vaccinees mount an equivalent magnitude of Spike-specific T cell response (both IFN-γ and IL-2 secretion) while higher levels of IFN-γ secretion were only detected in individuals who had an asymptomatic SARS-CoV-2 infection (Figure 5B). The latter observation is in line with previous analysis of asymptomatic and symptomatic SARS-CoV-2 infected individuals within 1 month of viral clearance where the former showed increased cytokine production with comparable frequencies of virus-specific T cells (5). Subtle qualitative differences in the Spike-specific T cell response were also observed in the different groups. Upon vaccination, Spike peptide induced IFN-γ and IL-2 secretion was comparable, but gradually diverged with time leading to higher levels of detectable IL-2 2-3 months post boost vaccination (Figure 5C). Symptomatic individuals also produced significantly more IL-2 than IFN-γ >6 months after viral clearance while this difference was less pronounced in asymptomatic individuals even at the latest time points tested (9-12 months post viral clearance) which did not reach statistical significance (Figure 5C). In individuals with previous SARS-CoV-1 infection, whole blood CRA also detected higher IL-2 secreting Spike-specific T cell responses 18 years after resolution of infection (Figure 5A). Hence, quantifying IL-2 secretion provides better sensitivity over IFN-γ in the detection of individuals with a long-term Spike-specific memory T cell response.

### Whole blood CRA detects the wide dynamic range and heterogeneous function of Spike-specific T cell responses in vaccinated individuals

In addition to evaluating the kinetics, quantitative and qualitative differences in the T cell response, the whole blood CRA also detected a wide range of Spike-specific T cell response in vaccinated individuals. Figure 6A shows the paired longitudinal samples of 27 vaccinees at ~14 and 90 days post boost vaccination. Secreted IFN-γ and IL-2 amounts in whole blood CRA differed between the two time points and among individuals. Interestingly, the quantity of cytokines detected two weeks after boost dose vaccination does not always predict the level of Spike-specific T cell response measurable at day 90. Some individuals exhibit >20-fold reduction at day 90 after boost vaccination, while in others, particularly for IL-2, the level was more stable (Figure 6). Indeed some individuals have a more pronounced decline of IFN-γ than IL-2 or vice versa as indicated in Figure 6B where the trajectory of the level of IFN-γ and IL-2 at day 14 and day 90 in each individual was plotted (Figure 6B).

**Figure 6.**
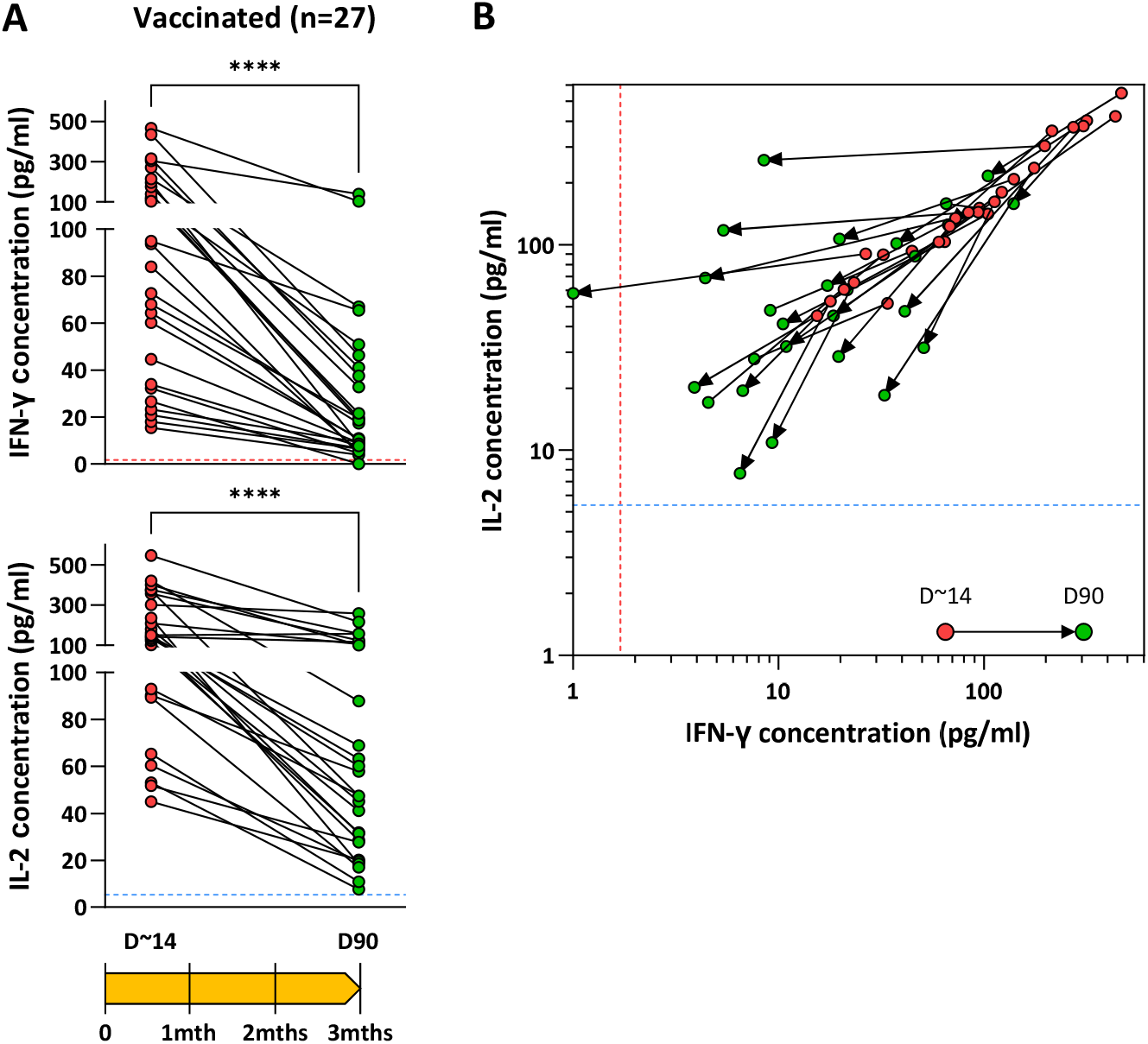
Heterogeneity of Spike-specific T cell response in vaccinated individuals. **A)** SARS-CoV-2 Spike-specific T cell response evaluated by SpG peptide pool stimulation of whole blood from vaccinated individuals (n=27) at 2 weeks (D~14; green circle) and 3 months (D90; red circle) after boost vaccination dose. Secreted IFN-γ and IL-2 concentrations detected are shown. **B)** Bivariate dot plots of secreted IFN-γ and IL-2 concentrations. Arrows connect paired individuals analyzed at D~14 and D90. Dashed lines denote the detection cut-off for the measured cytokines.

### Spike-specific T cell responses do not correlate with neutralizing antibody quantities

Lastly, since the quantification of serum neutralizing antibodies is the mainstay of SARS-CoV-2 humoral immunity assessment, we determined if there is a predictive relationship between the quantity of neutralizing antibodies and Spike-specific T cell responses. We performed linear regression analysis of the T cell response in vaccinated and convalescent asymptomatic and symptomatic COVID-19 patients quantified through whole blood CRA (both IFN-γ and IL-2) with the serum neutralizing antibody quantities assessed through RBD-hACE2 binding inhibition assay (Figure 7). No substantial correlations were observed in any of the analyzed groups. As also demonstrated previously in convalescent {PhD:2021bm} and in vaccinated individuals (26), serum neutralizing antibody quantities cannot predict the corresponding Spike-specific T cell responses in an individual and this further stresses how T cell response information from the whole blood CRA complements existing antibody assessments.

**Figure 7.**
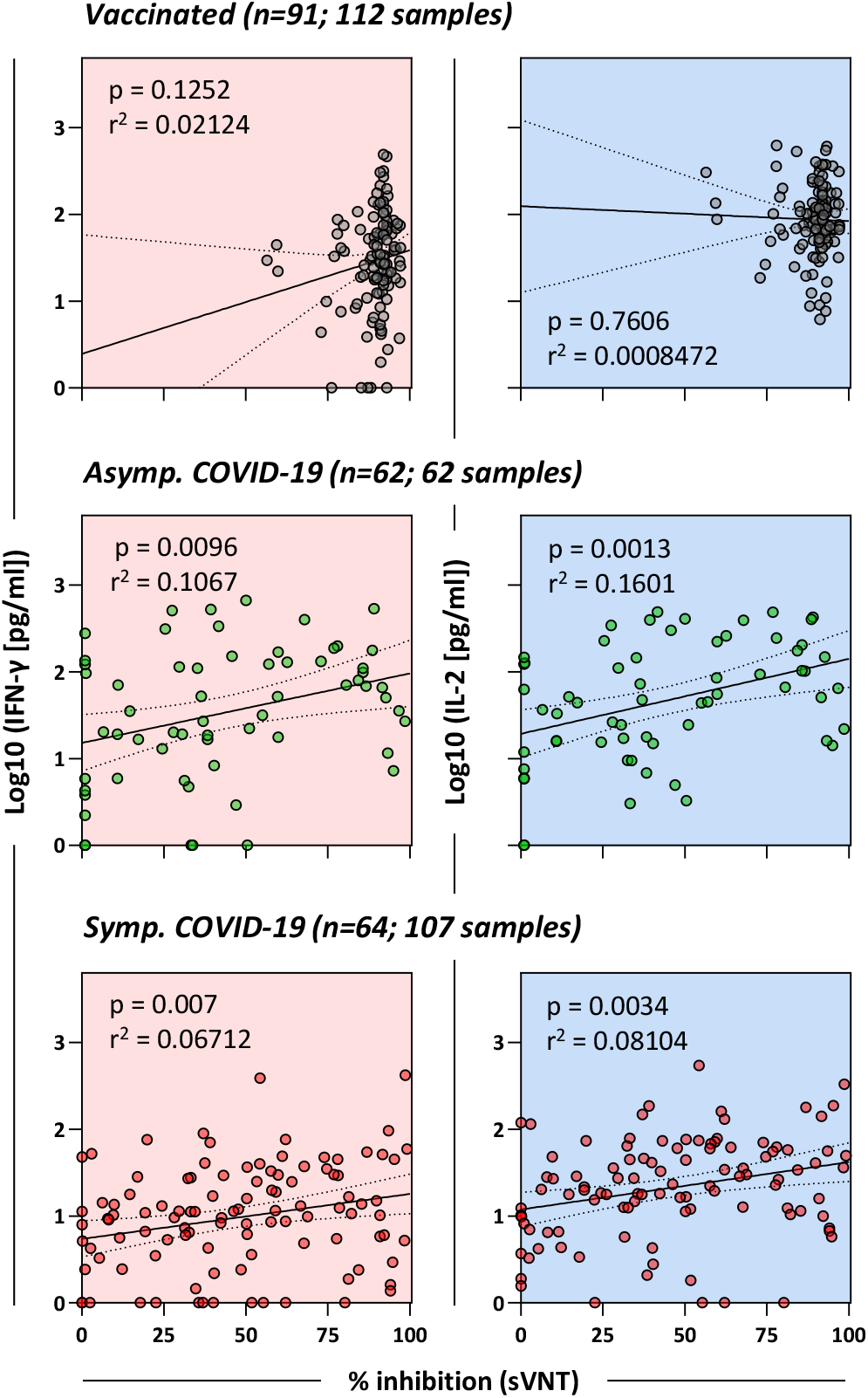
Spike-specific T cell responses do not correlate with the quantity of neutralizing antibodies in the serum. Linear regression analysis of the concentration of IFN-γ and IL-2 in SpG peptide pool stimulated whole blood and the corresponding SARS-CoV-2 neutralizing capacity of serum from vaccinated individuals (n=91; 112 samples), asymptomatic (n=62; 62 samples) and symptomatic (n=64; 107 samples) COVID-19 patients sampled at all timepoints available. Dotted lines denote the 95% confidence interval.

## Discussion

The complexity of virus-specific T cell characterization relegates their analysis to studies performed in selected laboratories accustomed with the complex methods of T cell analysis. The rapidity, simplicity and accuracy of the cytokine release assay (CRA) in whole blood can allow a routine measurement of SARS-CoV-2 T cells in large populations and thus help in understanding the role of antiviral T cells during the current COVID-19 pandemic. It is important to note that cytokine release in the stimulated whole blood does not detect the mere presence of T cells but also their functionality. This is an important feature of the assay that differentiates the CRA (and other assays like ELISPOT and AIM) from the recently developed test (T-Detect COVID, Adaptive Biotechnologies), which uses next generation sequencing of T cell receptors (TCR) to determine the presence/absence of cellular immunity to SARS-CoV-2 (27). We think that the ability to measure the wide dynamic range of functional Spike-specific T cells and not only their presence will be an important asset that will more precisely evaluate the protective ability of T cells after infection or vaccination.

In this work, by sequentially testing vaccinated and SARS-CoV-2 convalescent individuals, we show that IL-2 and IFN-γ quantification in whole blood measured Spike-specific T cell response with an accuracy equivalent to ELISPOT assays performed in freshly purified PBMC. Minor discrepancies between the magnitudes of T cells were detected only at early time points when CRA was able to detect a signal in the absence of ELISPOT results. Furthermore, analysis at 2 and 3 months after vaccination showed, on average, a better sensitivity of IL-2 over IFN-γ in the detection of subjects with Spike-specific T cell responses. The superior ability of IL-2 to detect long-term memory Spike-specific T cells was also supported by the analysis of SARS-CoV-2 convalescent individuals 12 months after infection and also 17 years after infection in the case of SARS-CoV-1 infected individuals. Interestingly, this difference in sensitivity is less pronounced in convalescent COVID-19 patients with asymptomatic infection. Though speculative at the moment, this lack of difference between IFN-γ and IL-2 secretion could reflect a better functionality of Spike-specific T cell response and hence plausibly contribute to the benign disease trajectory in these individuals.

The ability of CRA to measure the dynamic range of Spike-specific T cell responses allowed us to study a wide group of vaccinated and COVID-19 convalescent individuals. We observed that despite the homogeneous cohort of vaccinated adults (21-60 years of age, healthy and SARS-CoV-2 naive individuals), the Spike-specific T cell response was quantitatively different immediately after both the prime and vaccination doses (~14 days after vaccination) and after 3 months, similar to what was detected in a more heterogeneous population of COVID-19 convalescent individuals with mild or asymptomatic disease. In the vaccinated cohort, we first observed that half of the vaccinated individuals showed a peak of their Spike-specific T cell response ~14 days after the prime dose while half reached their peak response after the boost dose. These data were derived from a very limited sample size since we studied only 6 individuals at multiple time points after vaccination. However, the inability of the second vaccine dose to boost Spike-specific T (and antibody) responses in some individuals has been observed so far only after vaccination of SARS-CoV-2 convalescents and not, like in our study, in SARS-CoV-2 naïve. A possibility is that the pre-existing Spike-specific T cells primed by other coronaviruses could have influenced this different kinetic.

The analysis of a much larger cohort of vaccinated individuals (n=112) studied with less frequent sampling up to 3 months upon completion of vaccination further confirmed the heterogeneity of the magnitude of the induced Spike-specific cellular response. The quantity and the kinetics of decline of the Spike-specific cellular responses diverged among the vaccinated individuals with CRA IFN-γ and IL-2 concentrations that spans from 10-500pg/ml and 40-500pg/ml respectively. Interestingly, some individuals have a constant cytokine secretion level over the observation period while in others, the cytokines quantity drops precipitously. In addition, the longitudinal quantification of two cytokines further increased the heterogeneity of the vaccine-induced immunity in different individuals as IFN-γ and IL-2 quantities did not decrease in parallel in all the individuals. Some vaccinated individuals displayed a stable IL-2 production associated with a profound decrease of IFN-γ, while others showed exactly the opposite. Whether these differences can be attributed to the presence of different populations of effector/memory T cells or different ratios of CD4 and CD8 T cells and whether such differences have an impact on protection will need to be analyzed in a large clinical study. In addition, it will be important to continue monitoring the Spike-specific T cell response beyond the 3 months observation period to understand whether the Spike-specific T cell response induced by vaccines will behave like the one observed after natural infection. Importantly, the rapid cytokine assay was able to detect Spike-specific T cells in ~84% of SARS-CoV-2 individuals 1 year after infection and also in 8 out of 12 SARS-CoV-1 patients 18 years after infection.

Another important observation that further supports the concept of marked heterogeneity of the immune response induced by the BNT162b2 vaccination was the lack of correlation between the magnitude of humoral and cellular immunity. The substantial independence of different components of the immune system after the initial induction phase has been demonstrated in COVID-19 convalescents (28, 29) and can be explained by recent data showing that neutralizing antibodies can be produced without follicular T cell help (30). The CRA also reveals a similar profile both in COVID-19 convalescent and vaccinated individuals where an independence of the neutralizing antibody titers and the magnitude of cytokines secreted was detected. COVID-19 convalescents were studied over 1 year after infection and displayed different levels of neutralizing antibodies, while vaccinated individuals mostly displayed a consistently high neutralizing antibody titers with little variations among the tested individuals 3 months after vaccination. In other studies, antibody persistence was still observed up to 6 months post mRNA vaccination at the time of writing(31). We do not know whether antibodies induced by vaccination will show a similar rate of decline as observed after natural infection beyond the 6 months follow up till date, which can make the analysis of cellular immunity even more important. Our data at the moment show that three months after vaccination, the levels of neutralizing antibodies cannot be used as a surrogate of Spike-specific cellular immunity induced by vaccination.

This quantitative heterogeneity was however not mirrored by the regions of Spike protein targeted by T cells induced by BNT162b2 vaccination. We observed a substantial similarity of immunodominance among the different vaccinated individuals, with large part of the T cell response directed towards the Spike 2 chain and with an almost complete lack of T cell determinants within the N terminal region of S1 (region 501-705AA). A reduced presence of T cell epitopes in this region was already observed in SARS-CoV-2 convalescents (23). It will be interesting to test whether this documented profile of Spike-T cell specificity will also occur in individuals vaccinated with different products where subtle differences of codon usage, signal peptide and amino acid modifications (pre-fusion conformation stabilization and furin cleavage site mutations) were introduced(32) (33–35).

In conclusion, we show here that the rapid measurement of cytokine production in whole blood after peptide specific activation is a quick and simple assay that can reliably detect the wide dynamic range of functionally heterogeneous Spike-specific T cell response induced after vaccination or infection in different individuals. Even though T cells cannot prevent infection in the absence of antibodies, their pivotal role in the protection from disease severity has been shown in natural infection of normal (2) (3), oncological patients (7) and in vaccinated individuals (16). As such, since the quantity of Spike-specific T cells cannot be predicted by the simple measurement of antibodies, this higher throughput and simple assay represents a feasible approach to implement in routine testing to complement existing antibody measurements and thus help to define the correlates of protection necessary for the design of current vaccine strategies.

## Materials and Methods

### Subject details

Individuals who have recovered from SARS-CoV-2 infection (Asymptomatic: n=62; Symptomatic: n=68), vaccinated with BNT162b2 (n=112) or had SARS-CoV-1 infection 17 years ago (n=12) were enrolled in this work as part of the PROTECT study (National Healthcare Group Domain Specific Review Board, NHG DSRB ref. 2012/00917), Healthcare Worker Vaccination study (SingHealth Centralised Institutional Review Board, CIRB ref. 2021/2014), Novel Pathogens study (CIRB ref. 2018/3045) and SARS Recall study (NHG DSRB ref. 2020/00091). All participants provided written informed consent. Vaccinated individuals were between 21-60 years of age, healthy and SARS-CoV-2 infection naïve. Whole blood and serum samples were collected at the indicated intervals for serological and T cell response analysis.

### Peptides

15-mer peptides that are overlapping by 10 amino acids (AA) spanning the entire SARS-CoV-2 Spike protein (GISAID EPI_ISL_410713) were synthesized (Genscript) and pooled into 7 pools of approximately 40 peptides in each pool (Supplementary Table 1). 55 Spike peptides covering the immunogenic regions of the SARS-CoV-2 Spike protein that represents 40.5% of the whole Spike protein forms the SpG peptide pool as described previously (5).

### Cytokine release assay (CRA) from whole peripheral blood stimulated with SARS-CoV-2 Spike peptide pools

320 μl of whole blood drawn on the same day were mixed with 80 μl RPMI and stimulated with the indicated SARS-CoV-2 Spike peptide pools (Supplementary Table 1) at 2 μg/ml or with DMSO as a control. After 16 hours of culture, the culture supernatant (plasma) was collected and stored at −80°C. Cytokine concentrations in the plasma were quantified using an Ella machine with microfluidic multiplex cartridges measuring IFN-γ and IL-2 following the manufacturer’s instructions (ProteinSimple). The level of cytokines present in the plasma of DMSO controls was subtracted from the corresponding peptide pool stimulated samples. The positivity threshold was set at 10x times the lower limit of quantification of each cytokine (IFN-γ = 1.7pg/ml; IL-2 = 5.4pg/ml) after DMSO background subtraction.

### Peripheral blood mononuclear cell isolation

Peripheral blood of all individuals was collected in heparin containing tubes and peripheral blood mononuclear cells from all collected blood samples were isolated by Ficoll-Paque density gradient centrifugation.

### SARS-CoV-2 Spike-specific T cell quantification

The frequency of SARS-CoV-2 Spike-specific T cells was quantified as described previously (5).Briefly, freshly isolated or cryopreserved PBMC (as indicated) were stimulated with SpG peptide pool in an IFN-γ ELISPOT assay. ELISPOT plates (Millipore) were coated with human IFN-γ antibody overnight at 4°C. 400,000 PBMC were seeded per well and stimulated for 18h with the SpG peptide pool at 2 μg/ml. The plates were then incubated with human biotinylated IFN-γ detection antibody, followed by Streptavidin-AP and developed using the KPL BCIP/NBT Phosphatase Substrate. To quantify positive peptide-specific responses, 2× mean spots of the unstimulated wells were subtracted from the peptide-stimulated wells, and the results expressed as spot forming cells (SFC)/10^6^ PBMC. Results were excluded if negative control wells had >30 SFC/10^6^ PBMC or if positive control wells (PMA/Ionomycin) were negative.

### RBD-hACE2 binding inhibition assay

Antibodies inhibiting virus binding to host cell was measured using a commercial RBD-human angiotensin-converting enzyme 2 (hACE2) binding inhibition assay called cPASS™ (GenScript). As per manufacturer’s instructions, serum was diluted 1:10 in the kit sample buffer and mixed 1:1 with HRP-conjugated RBD and incubated for 30 mins at 37°C. RBD-antibody mixtures were then transferred and incubated for 15 mins at 37°C in enzyme-linked immunosorbent assay (ELISA) plates coated with recombinant hACE2 receptor. Following incubation, plates were washed with the kit wash solution, incubated with TMB substrate for 15 mins and reaction stopped with stop solution. Absorbance was measured at OD450 nm. Percent inhibition of RBD-hACE2 binding was computed using the following equation: % inhibition=(1-[(OD of serum+RBD)/(OD of negative control+RBD)])×100. As described by the cPASS™ kit, a cutoff of 20% and above was used to determine positive RBD-hACE2 inhibition.

### Activation induced cell marker assay

Cryopreserved PBMCs were thawed and stimulated for 24 hr at 37^°^C with the SpG peptide pool (2μg/ml) in AIM-V media supplemented with 2% AB serum. Cells were then stained with the Fixable Near-IR Live/Dead fixable cell stain kit (Invitrogen) followed with surface markers as previously described (16): anti-CD3, anti-CD4, anti-CD8, anti-CD69, anti-CD134 (OX40) and anti-CD137 (4-1BB). All samples were acquired on a BD-LSR II Analyzer (BD) and analyzed with FlowJo software (BD). Gating strategy is shown in Supplementary Figure 2.

### Statistical analysis

All statistical analyses were performed using GraphPad Prism v9. Where applicable, the statistical tests used and the definition of centre were indicated in the figure legends. Statistical significance was defined as having a P-value of less than 0.05. In all instances, “n” refers to the number of patients analysed.

## Author contributions

A.T. Tan, N. Le Bert and A. Bertoletti designed the experiments; W.N. Chia, R. Alwiss, E.E. Ooi and L-F. Wang performed and analyzed the antibody experiments; J.M.E. Lim, K. Kunasegaran, A. Chia, M.D.C. Qui and N. Tan performed all other experiments and analyzed the data; A.T. Tan, N. Le Bert and A. Bertoletti analyzed and interpreted all the data; A.T. Tan, N. Le Bert and A. Bertoletti prepared the figures and wrote the paper; D. Ying, M.I. Cheng, B. Young, L.Y. Hsu, J.GH. Low and D.C. Lye recruited all the COVID-19 patients, SARS-recall patients and the vaccinees and provided all clinical samples and data; A. Bertoletti designed and coordinated the study.

## Acknowledgments

We would like to thank all clinical and nursing staff who provided care for the patients; all clinical trial coordinators and staff in Singapore Infectious Disease Clinical Research Network and Infectious Disease Research and Training Office, National Centre for Infectious Diseases and Singapore General Hospital for their invaluable assistance in coordinating patient recruitment.

---

**Supplementary Figure 1.**
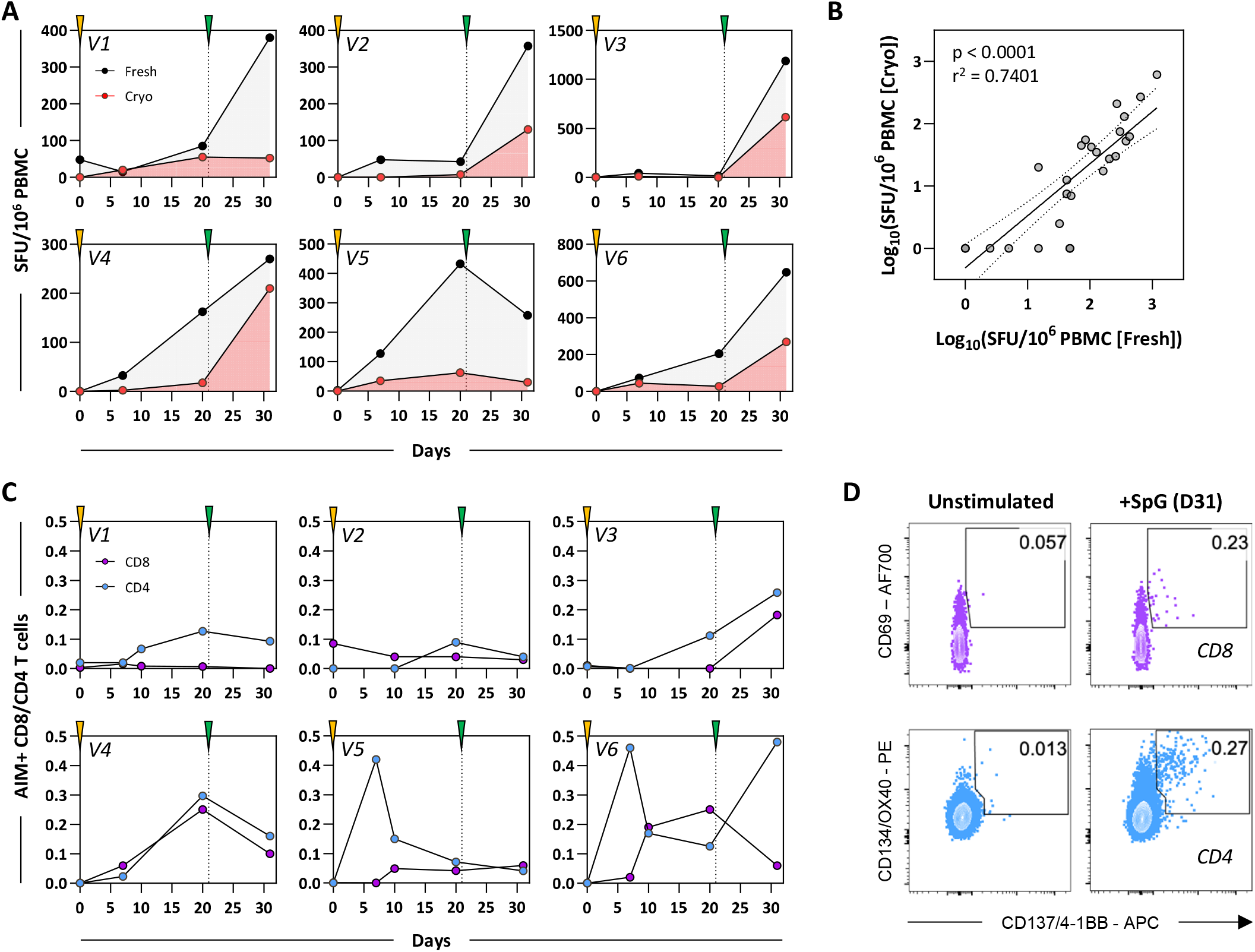
Longitudinal response dynamics of ELISPOT assay performed using fresh and cryopreserved PBMCs. **A)** Standard IFN-γ ELISPOT assays using SpG peptide pool were performed in parallel using freshly isolated (black) or cryopreserved PBMCs (red) obtained from vaccinated individuals at different time points post vaccination (n=6; 24 samples). **B)** Linear regression analysis of IFN-γ SFU/10^6^ PBMCs quantified using freshly isolated or cryopreserved PBMCs shows a strong positive correlation. Dotted lines denote the 95% confidence interval. **C)** The frequency of Spike-specific CD4 and CD8 T cells 10 days after the second dose of BNT162b2 were quantified from cryopreserved PBMC samples through the induction of activation markers (CD69, 4-1BB and/or OX40) after SpG peptide pool stimulation (n=6; 27 samples). **D)** Flow cytometry bivariate dot plots showing AIM+ CD8 and CD4 T cells frequencies after SpG peptide pool stimulation in a representative vaccinee 10 days after the boost dose.

**Supplementary Figure 2.**
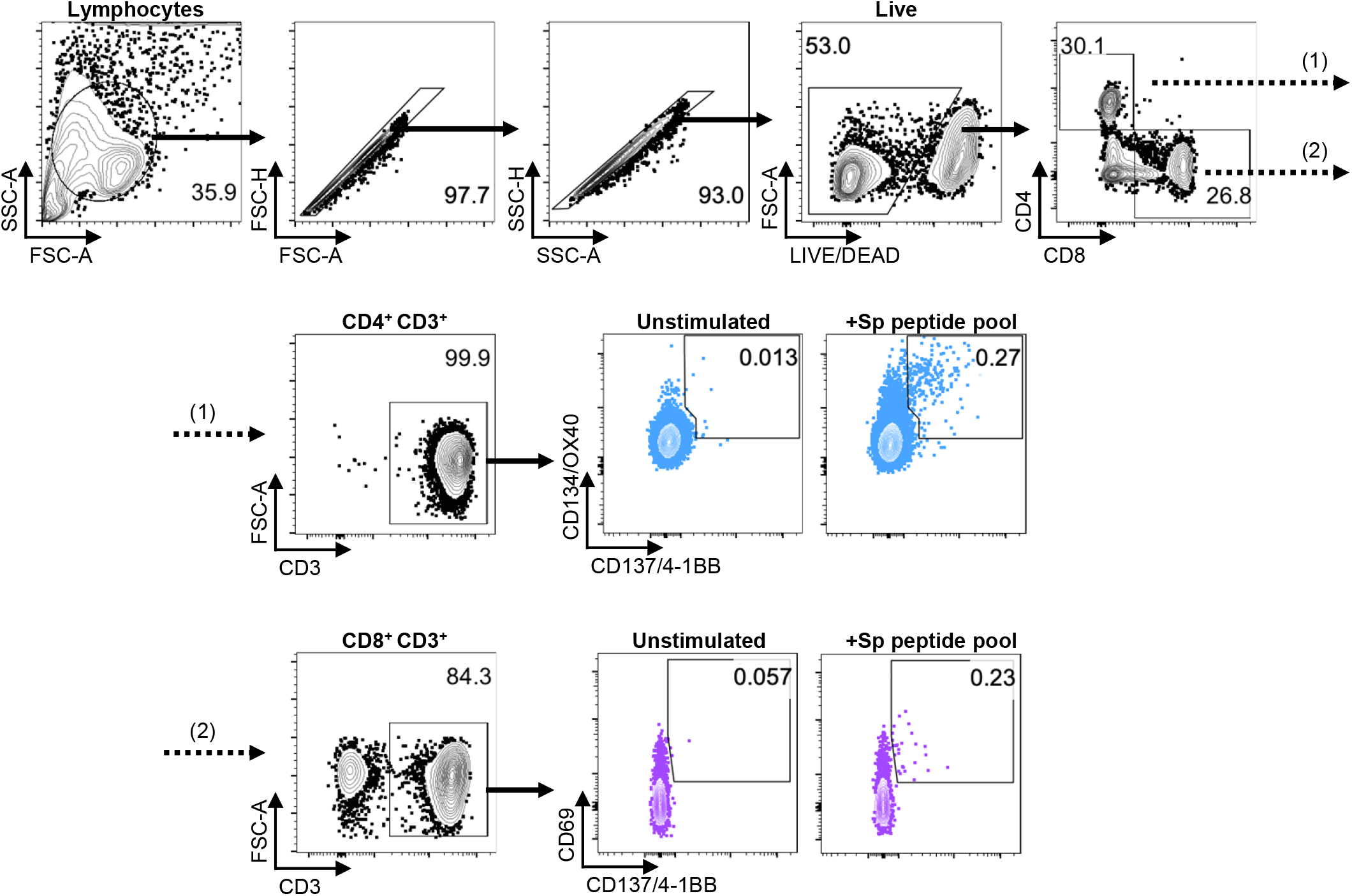
Representative gating strategy for AIM+ CD4+ (CD134/OX40+ CD137/4-1BB+; blue) and CD8+ (CD69+ CD137/4-1BB+; purple) T cells in PBMCs with and without peptide stimulation. Frequencies of gated populations relative to parent are affixed into the contour plots.

**Supplementary Figure 3.**
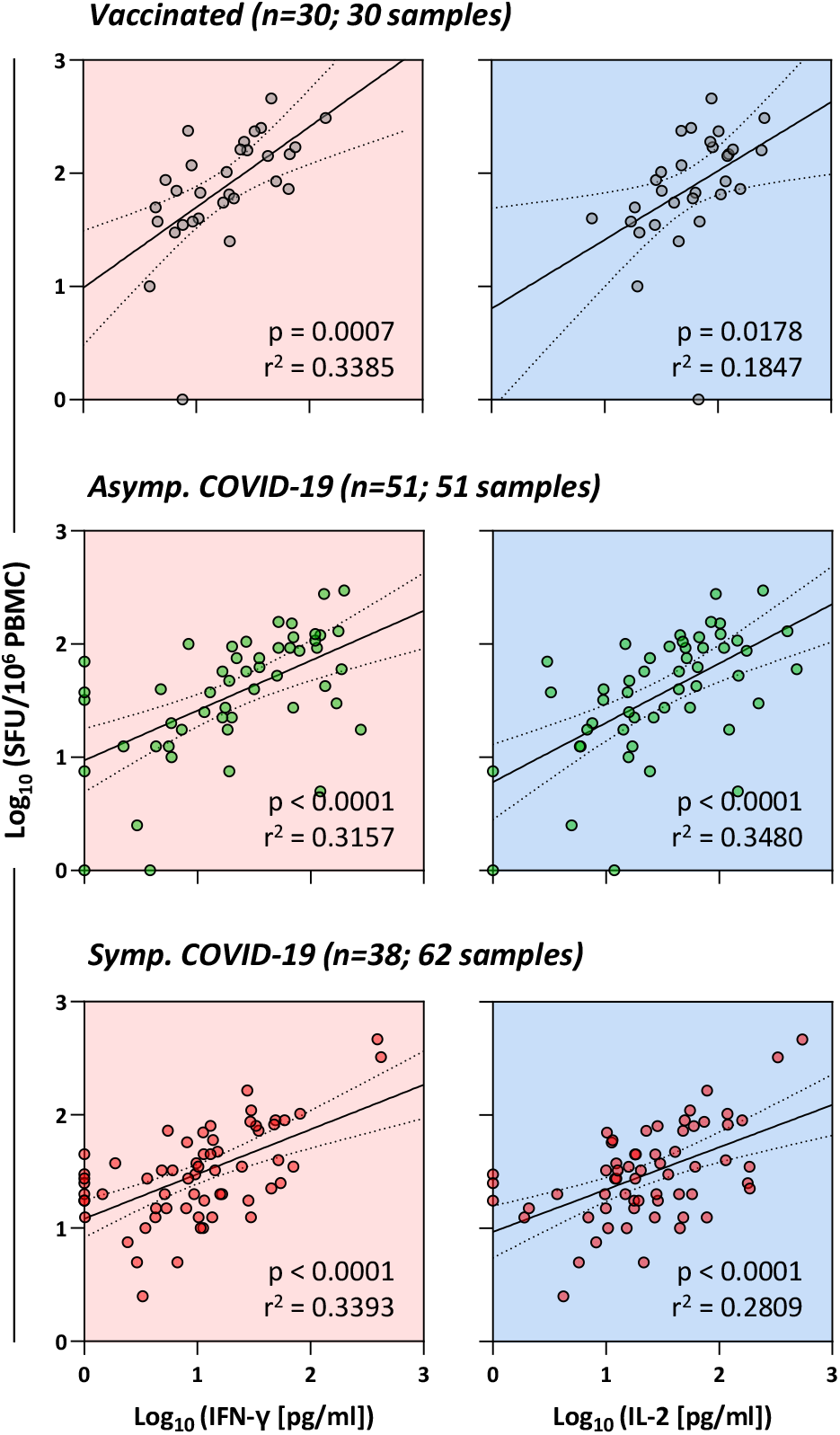
Whole blood CRA quantifies Spike-specific memory T cell responses ≥3 months post-boost vaccination dose or viral clearance. Linear regression analysis of the concentration of IFN-γ and IL-2 in SpG peptide pool stimulated whole blood and the corresponding frequency of Spike-specific T cells quantified by IFN-γ ELISPOT in vaccinated individuals (grey; n=30; 30 samples), asymptomatic (green; n=51; 51 samples) and symptomatic (red; n=38; 62 samples) COVID-19 patients sampled ≥3 months post-boost vaccination dose or viral clearance. Dotted lines denote the 95% confidence interval.

**Supplementary Figure 4.**
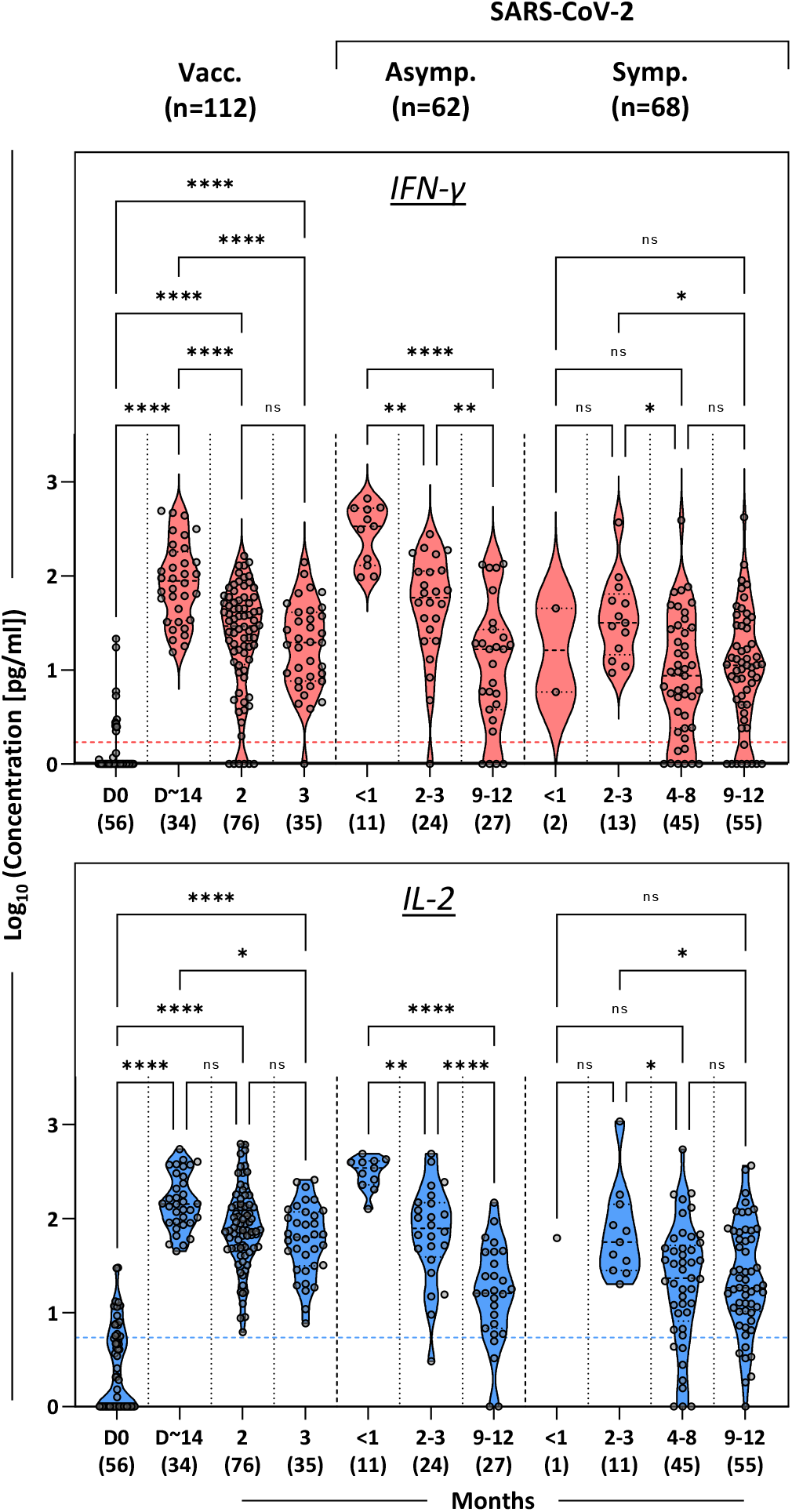
SARS-CoV-2 spike specific T cells in the whole blood of vaccinated and infected individuals. **A)** SARS-CoV-2 Spike-specific T cell response in vaccinated individuals (n=112; 201 samples), asymptomatic (n=62; 62 samples) and symptomatic (n=68; 115 samples) COVID-19 patients were longitudinally quantified by measuring IFN-γ (red) and IL-2 (blue) secretion in whole blood after SpG peptide pool stimulation. Dashed lines denote the detection cut-off for the measured cytokines. The number of samples analysed at each time point were indicated in parentheses. Significant differences in each group were analysed by one-way ANOVA and the adjusted p-value (adjusted for multiple comparison) are shown. ns = not significant P>0.05; * = P≤0.05; ** = P≤0.01; *** = P≤0.001; **** = P≤0.0001.

**Supplementary Table 1.**
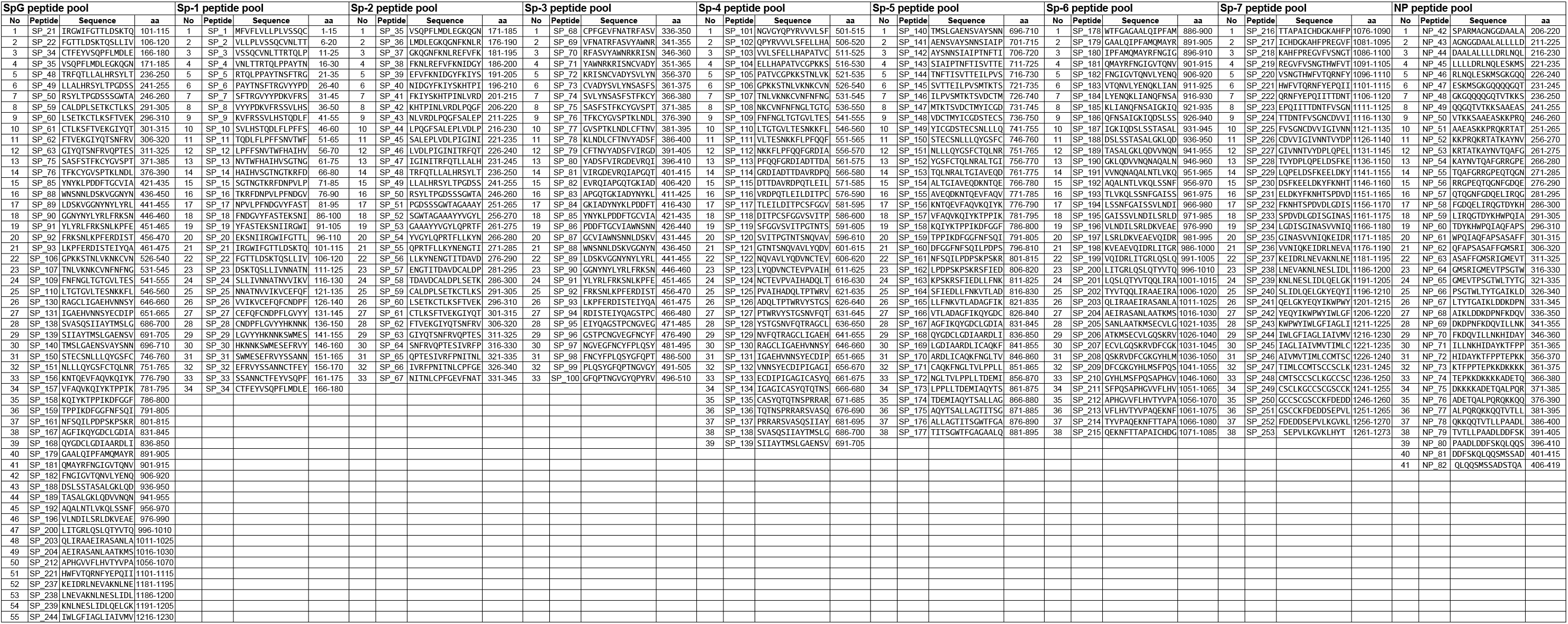
Details of the peptides found in the SpG pool, and the overlapping peptide pools covering the entire SARS-CoV-2 Spike protein and the C-terminal half of the nucleoprotein.

## References

1. McMahan K et al. Correlates of protection against SARS-CoV-2 in rhesus macaques. Nature 2021;590(7847):630–634.

2. Rydyznski Moderbacher C et al. Antigen-Specific Adaptive Immunity to SARS-CoV-2 in Acute COVID-19 and Associations with Age and Disease Severity. Cell 2020;183(4):996–1012.e19.

3. Tan AT et al. Early induction of functional SARS-CoV-2-specific T cells associates with rapid viral clearance and mild disease in COVID-19 patients. Cell Reports 2021;53:108728–13.

4. Sekine T et al. Robust T Cell Immunity in Convalescent Individuals with Asymptomatic or Mild COVID-19. Cell 2020;183(1):158–168.e14.

5. Le Bert N et al. Highly functional virus-specific cellular immune response in asymptomatic SARS-CoV-2 infection. J Exp Med 2021;218(5). doi:10.1084/jem.20202617

6. Rodda LB et al. Functional SARS-CoV-2-Specific Immune Memory Persists after Mild COVID-19. Cell 2021;184(1):169–183.e17.

7. Bange EM et al. CD8+ T cells contribute to survival in patients with COVID-19 and hematologic cancer. Nat Med 2021; doi:10.1038/s41591-021-01386-7.

8. Walsh EE et al. Safety and Immunogenicity of Two RNA-Based Covid-19 Vaccine Candidates. N Engl J Med 2020;:NEJMoa2027906–13.

9. Jackson LA et al. An mRNA Vaccine against SARS-CoV-2 — Preliminary Report. N Engl J Med 2020;383(20):1920–1931.

10. Sahin U et al. COVID-19 vaccine BNT162b1 elicits human antibody and TH1 T cell responses. Nature 2020;586(7830):594–599.

11. Khoury DS et al. Neutralizing antibody levels are highly predictive of immune protection from symptomatic SARS-CoV-2 infection. Nat Med 2021;:1–7.

12. Chia WN et al. Dynamics of SARS-CoV-2 neutralising antibody responses and duration of immunity: a longitudinal study. Lancet Microbe 2021;2(6):e240–e249.

13. Sahin U et al. BNT162b2 vaccine induces neutralizing antibodies and poly-specific T cells in humans. Nature [published online ahead of print: May 27, 2021]; doi:10.1038/s41586-021-03653-6

14. Bonifacius A et al. COVID-19 immune signatures reveal stable antiviral T cell function despite declining humoral responses. Immunity 2021;54(2):340–354.e6.

15. Sherina N et al. Persistence of SARS-CoV-2-specific B and T cell responses in convalescent COVID-19 patients 6-8 months after the infection. Med (N Y) 2021;2(3):281–295.e4.

16. Kalimuddin S et al. Early T cell and binding antibody responses are associated with Covid-19 RNA vaccine efficacy onset. Med (N Y) [published online ahead of print: April 8, 2021]; doi:10.1016/j.medj.2021.04.003

17. Wyllie D et al. SARS-CoV-2 responsive T cell numbers are associated with protection from COVID-19: A prospective cohort study in keyworkers. medRxiv 2020;3(5):e2010182–24.

18. Goletti D et al. Selected RD1 peptides for active tuberculosis diagnosis: comparison of a gamma interferon whole-blood enzyme-linked immunosorbent assay and an enzyme-linked immunospot assay. Clin Diagn Lab Immunol 2005;12(11):1311–1316.

19. Petrone L et al. A whole blood test to measure SARS-CoV-2-specific response in COVID-19 patients. Clin Microbiol Infect 2021;27(2):286.e7–286.e13.

20. Murugesan K et al. Interferon-gamma release assay for accurate detection of SARS-CoV-2 T cell response. Clin. Infect. Dis. [published online ahead of print: October 9, 2020]; doi:10.1093/cid/ciaa1537

21. Li J et al. Safety and immunogenicity of the SARS-CoV-2 BNT162b1 mRNA vaccine in younger and older Chinese adults: a randomized, placebo-controlled, double-blind phase 1 study. Nat Med 2021;:1–9.

22. Ford T et al. Cryopreservation-related loss of antigen-specific IFNγ producing CD4+ T-cells can skew immunogenicity data in vaccine trials: Lessons from a malaria vaccine trial substudy. Vaccine 2017;35(15):1898–1906.

23. Tarke A et al. Comprehensive analysis of T cell immunodominance and immunoprevalence of SARS-CoV-2 epitopes in COVID-19 cases. Cell Reports Medicine 2021;181(D1):100204.

24. Woldemeskel BA, Garliss CC, Blankson JN. SARS-CoV-2 mRNA vaccines induce broad CD4+ T cell responses that recognize SARS-CoV-2 variants and HCoV-NL63. J. Clin. Invest. 2021;131(10). doi:10.1172/JCI149335

25. Le Bert N et al. SARS-CoV-2-specific T cell immunity in cases of COVID-19 and SARS, and uninfected controls. Nature 2020;584(7821):457–462.

26. Reynolds CJ et al. Prior SARS-CoV-2 infection rescues B and T cell responses to variants after first vaccine dose. Science [published online ahead of print: April 30, 2021]; doi:10.1126/science.abh1282

27. Snyder TM et al. Magnitude and Dynamics of the T-Cell Response to SARS-CoV-2 Infection at Both Individual and Population Levels. medRxiv 2020;:1–29.

28. Reynolds CJ et al. Discordant neutralizing antibody and T cell responses in asymptomatic and mild SARS-CoV-2 infection. Science Immunology 2020;5(54). doi:10.1126/sciimmunol.abf3698

29. Dan JM et al. Immunological memory to SARS-CoV-2 assessed for up to 8 months after infection. Science 2021;371(6529). doi:10.1126/science.abf4063

30. Chen JS et al. High-affinity, neutralizing antibodies to SARS-CoV-2 can be made in the absence of T follicular helper cells. bioRxiv 2021;:2021.06.10.447982.

31. Doria-Rose N et al. Antibody Persistence through 6 Months after the Second Dose of mRNA-1273 Vaccine for Covid-19. N Engl J Med 2021;384(23):2259–2261.

32. van Doremalen N et al. ChAdOx1 nCoV-19 vaccine prevents SARS-CoV-2 pneumonia in rhesus macaques. Nature 2020;586(7830):578–582.

33. Corbett KS et al. SARS-CoV-2 mRNA vaccine design enabled by prototype pathogen preparedness. Nature 2020;586(7830):567–571.

34. Bos R et al. Ad26 vector-based COVID-19 vaccine encoding a prefusion-stabilized SARS-CoV-2 Spike immunogen induces potent humoral and cellular immune responses. NPJ Vaccines 2020;5(1):91–11.

35. Vogel AB et al. BNT162b vaccines protect rhesus macaques from SARS-CoV-2. Nature 2021;592(7853):283–289.

